# Structural color in *Junonia* butterflies evolves by tuning scale lamina thickness

**DOI:** 10.1101/584532

**Authors:** Rachel C. Thayer, Frances I. Allen, Nipam H. Patel

## Abstract

In diverse organisms, nanostructures that coherently scatter light create structural color, but how such structures are built remains mysterious. We investigate the evolution and genetic regulation of butterfly scale laminae, which are simple photonic nanostructures. In a lineage of buckeye butterflies artificially selected for blue wing color, we found that thickened laminae caused a color shift from brown to blue. Deletion of the *optix* wing patterning gene also altered color via lamina thickening, revealing shared genetic regulation of pigments and lamina thickness. Finally, we show how lamina thickness variation contributes to the color diversity that distinguishes sexes and species throughout the genus *Junonia*. Thus, quantitatively tuning one dimension of scale architecture facilitates both the microevolution and macroevolution of a broad spectrum of hues. Because the lamina is an intrinsic component of typical butterfly scales, our findings suggest that tuning lamina thickness is a readily accessible mechanism to create structural color across the Lepidoptera.

## Introduction

Structural colors are both visually delightful and abundant in nature. Organisms deploy structural colors to display hues for which they lack pigments (frequently blues and greens), to create specific optical effects such as iridescence or light polarization, and to mediate ecological interactions, including intraspecific signaling and camouflage. Unlike pigmentary color, which is caused by molecules that selectively absorb certain wavelengths of light, structural colors result from the constructive and destructive interference of light as it interacts with nanoscale, precisely-shaped physical structures that are made of a high refractive index material (e.g. keratin, chitin, or cellulose).

Despite the clear importance of structural color for living systems, the biological production of structural colors has long eluded characterization [1]. Many experimental techniques depend on harnessing variation to dissect biological processes, but photonic structures are so small that quantitatively measuring variation in their dimensions is technically demanding, especially for high-throughput sampling, detecting subtle variation that may segregate within populations, or analyzing over developmental time *in vivo*. The color itself is easier to quantify, but has limited utility as a proxy for nanostructural dimensions, since structural colors and pigments often co-occur and covary. While recent studies [2–4] have made early headway toward describing genetic regulation of structural colors, much work remains to decipher the evolutionary, developmental, and genetic bases of structural coloration, and lab-tractable systems with intraspecific variation in structural coloration are needed. We present a promising system, the butterfly genus *Junonia*, with extensive variation in a simple structural color, and show how structural simplicity is a tactical advantage when seeking to unravel mechanisms for the biological production of nanostructures.

In butterflies, photonic nanostructures occur within the architecture of scales. Scales are the fundamental coloration unit on butterfly wings and have a Bauplan consisting of a grid of ridges and ribs, supported by a lower lamina that is a simple plane (Fig. 1A). Scales are composed of chitin and may also have embedded pigments. Intricate architecture and a high refractive index make scales a pliable substrate for photonic innovations, and indeed scales have been evolutionarily elaborated in many ways for impressive optical effects [5]. Even the simplest butterfly scales can produce structural color, via the lower lamina acting as a thin film reflector. Thin films are the simplest photonic structure and consist of a layer of high refractive index material, on the order of hundreds of nanometers thick, surrounded by a material with a contrasting refractive index, i.e. air (Fig. 1B). Light is reflected from each surface of the film, and these two reflections interfere with each other. If the two reflections remain in phase, which depends on the extra distance traveled through the film and the wavelength, then they interfere constructively to produce observable color [6,7]. Conversely, wavelengths (colors) that undergo destructive interference have decreased brightness.

**Figure 1:**
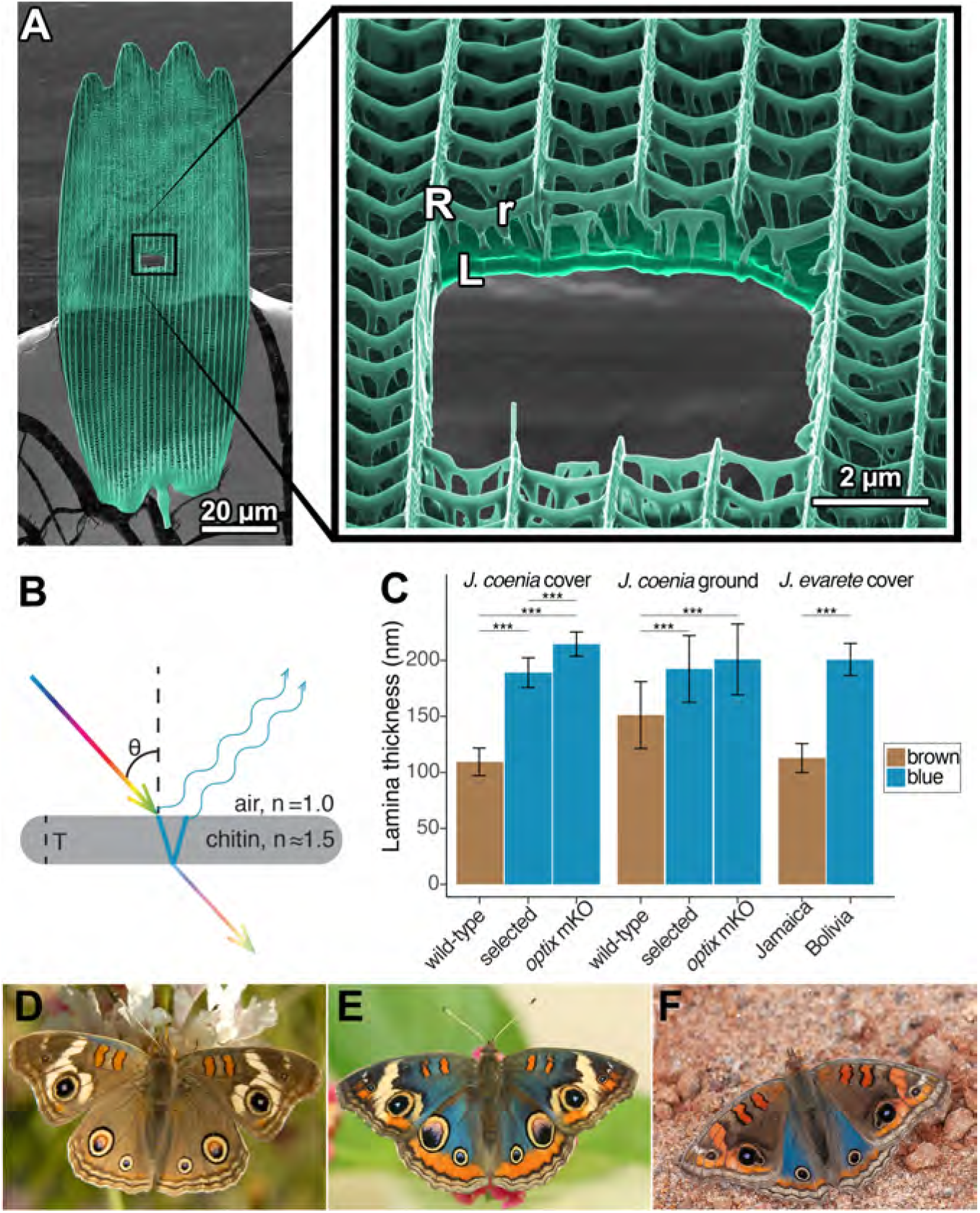
The lamina of a typical butterfly scale functions as a thin film reflector. *(A)* Colorized helium ion micrograph of a nymphaline scale, with a window milled using a gallium focused ion beam. Inset at higher magnification, with labels for general architectural components of a scale (R = ridges, r = cross-ribs, L = lamina). *(B)* Diagram of reflection and refraction in a chitin thin film. White light enters, reflections are produced at each surface of the film, and reflections of select wavelengths remain in phase as a function of film thickness (T). *(C)* Experimental disruptions of wing color are associated with altered lamina thickness. In *J. coenia*, artificial selection for blue color increased lamina thickness in both cover and ground scales. In *optix* mosaic knockout mutants, certain wing regions have similar thickness increases. This trend recapitulates natural variation in *J. evarete*, where blue butterflies have thick laminae relative to brown individuals. (*** = p < 1×10^−7^) Bars show mean thickness, with error bars of one standard deviation. *(D)* Wild-type *J. coenia. (E)* Blue artificially selected *J. coenia.* Image by Edith Smith. *(F) J. evarete*.

While it is known that the thickness of the lower lamina is one parameter that controls structural color wavelength [8] and that thickness can respond to artificial selection in the laboratory [9], it is not known how general this mechanism is in natural evolution. It is also unknown how lamina structural colors are genetically regulated and whether any recognized butterfly wing patterning genes regulate lamina thickness. Here, we use mutants with deletions in the *optix* wing patterning gene, artificial selection on wing color, and genus-wide wing color variation to test the role of lamina thickness in generating butterfly color. We show that butterflies in the genus *Junonia* thoroughly exploit the relationship between film thickness and color, using the thin films necessarily present in their scales to produce a broad spectrum of hues by tuning lamina thickness. These lamina colors work in tandem with pigments to define the wing pattern elements that distinguish populations, sexes, and species, indicating that the ability to vary lamina thickness has been an important microevolutionary and macroevolutionary tool in this group, and likely in butterflies more broadly.

## Results

### Artificial selection for blue wing color increases lamina thickness

Here we describe a novel instance of rapid, artificially selected color shift from brown to blue wing color in *J. coenia* buckeye butterflies (Fig 1D-E) and identify the structural changes that enabled the color shift. Edith Smith, a private butterfly breeder, began selectively mating buckeyes with a few blue scales on the costal margin of the dorsal forewing (E. Smith, personal communication, Sep. 2014). After five months of selective breeding, blue spread to the dorsal hindwing of some individuals. By eight months, there was a noticeable increase in blue surface area, and within roughly 12 months (on the order of 12 generations), most butterflies in the breeding colony were visibly blue over the majority of their dorsal wing surface. On the forewing, areas proximal to M1 were visibly blue, except the discal bars (Fig. S1). On the hindwing, blue shift did not include the distal-most wing pattern elements, i.e. EI-EIII and eyespots. At its strongest, the phenotype may include blue scales cupping the posterior forewing eyespot and/or a blue sheen in all distal elements of the forewing. Smith maintained the blue colony for several years, introgressing a few progeny from crosses to wild-caught buckeyes about once per year to maintain genetic diversity. Over time, she noted the emergence of a variety of short-wavelength colors, ranging from purple to green. Two years after focused selection, she estimated that the population was 85% blue, 8% green, 2% purple, and 5% brown. Like many familiar examples of human selection (e.g. domesticated animals, crop plants), outcomes are informative even without complete experimental documentation of the selective process [10,11]. These selected blue buckeyes provide a previously unexploited opportunity to study structural color. They demonstrate rapid and extensive evolutionary color change, and are a stark contrast to wild-type brown populations with which they are still interfertile. Conveniently, the artificially selected taxon, *J. coenia*, is a recognized model species for butterfly developmental genetics [12,13]. The selected blue individuals resemble naturally evolved color variants in the sister species, *J. evarete* (Fig. 1F), and offer a useful comparison to a previously reported artificial selection experiment in butterflies [9].

To pinpoint the cause of blueness in artificially selected butterflies, we characterized cover scales from the dorsal hindwing (Fig. 2A-D). Butterfly wings have two classes of scales arranged in alternating rows that form two layers: superficial cover scales and underlying ground scales. Cover and ground scales frequently have contrasting size, shape, and color, and their juxtaposition can be important for wing color [8]. When isolated and laid in the abwing orientation they occupy on the wing, cover scales were blue (Fig. 2B). However, when flipped over and viewed in adwing orientation, which exposes only the lower lamina, scales appeared more brightly blue and iridescence was more apparent (Fig. 2B’, 2D). We tested whether the blue was structural rather than pigment-based by immersing the full scale in oil with a refractive index matched to that of chitin (Fig. 2B’’’). Index-matching eliminates the possibility of reflection and structural color, leaving only pigment-based coloration. We measured the scale’s absorption spectrum under these conditions (Fig. 3A), which revealed that blue scales did have some pigment, presumably a brown ommochrome [14], but this pigment cannot account for blueness. The pigment was located in the scale ridges (Fig. S2 B). Lepidopteran structural colors may occur in the lamina, lumen, ridges, or cross-ribs. To isolate which of these features had the nanostructure responsible for blue structural color, we dissected the scales (Fig. 2B”, Fig. S2 A). After removing all other scale components, we found that the bare lower lamina was sufficient for blue structural color. We also examined regions with all scale components except the lamina and found that these pieces of lamina-less scale were not blue (Fig. S2 C). We thus focused on investigating nanostructure in the lamina. To discern between a single or multilayer lamina and take precise measurements, we cross-sectioned the lamina and viewed it with Helium Ion Microscopy (HIM) (Fig. 2C). HIM imaging indicated the lower lamina was a simple monolayer of chitin with a thickness of 187 ± 13 nm (SD, Fig. 1C), which is a reasonable thickness to reflect blue as a dielectric thin film [8].

**Figure 2:**
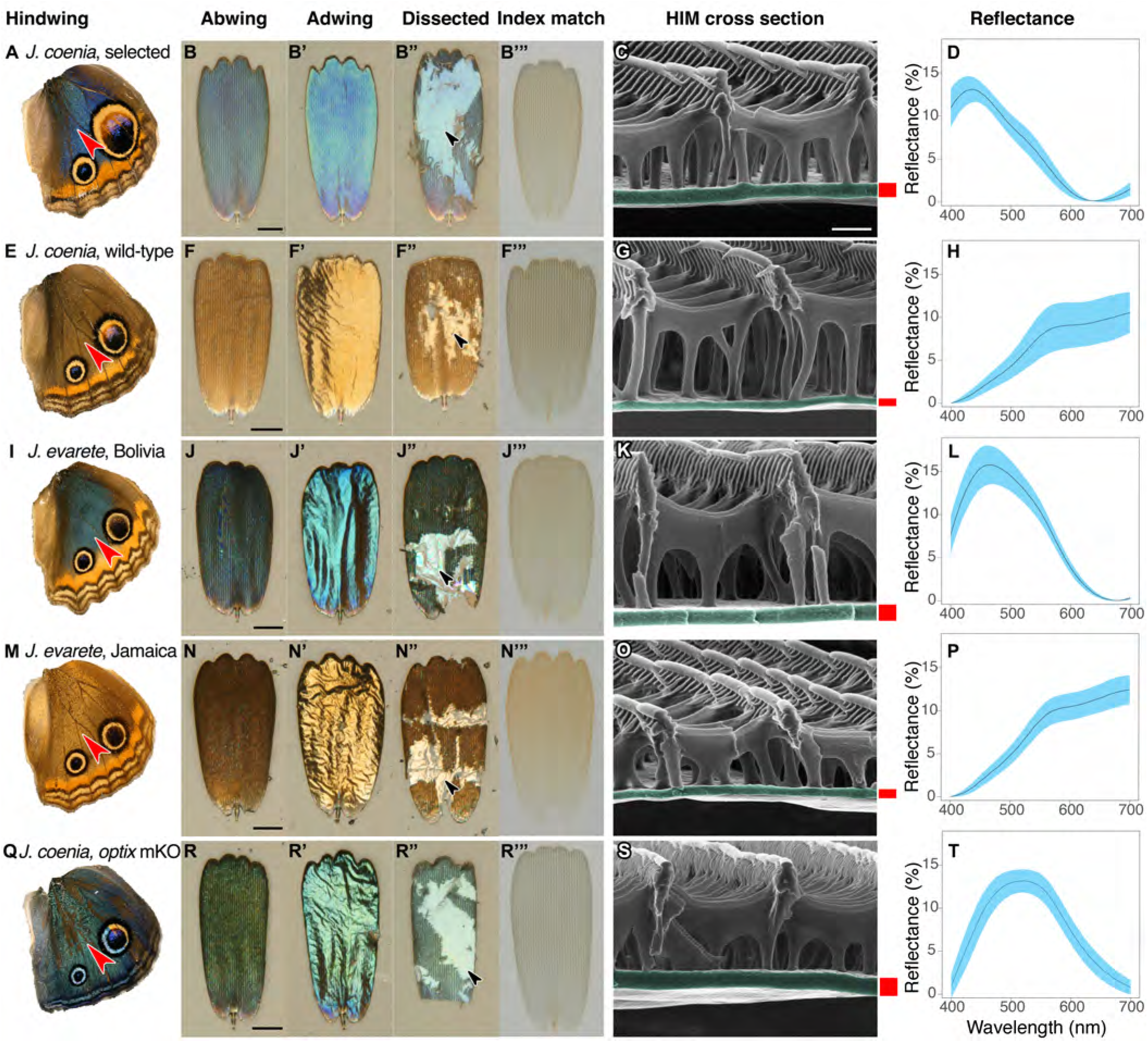
Structure and color of *Junonia* cover scales. *(A-D)* Artificially selected blue *J. coenia*. *(E-H)* Wild-type *J. coenia*. *(I-L) J. evarete*, blue male from Bolivia. *(M–P) J. evarete*, brown male from Jamaica. *(Q-T) optix* mosaic knockout mutant (mKO) in *J. coenia*. *(A,E,I,M,Q)* Dorsal hindwing, red arrowhead indicates the characterized scale’s location. *(B,F,J,N,R)* Scale in the orientation it would occupy on the wing, showing the abwing surface of the cover scale. Black scale bars are 25 µm. *(B’,F’,J’,N’,R’)* Adwing surface of cover scale, showing the underside of the lamina. *(B”,F”,J”,N”,R”)* Dissected scale with arrow showing regions where all ridges and ribs are removed to expose the bare lamina. The lamina is sufficient to create iridescent blue and gold structural colors. *(B”’,F”’,J”,N”’,R”’)* Scale immersed in fluid with a refractive index matched to chitin, thus eliminating reflection to show only pigmentary color. Blue and brown scales have comparable amounts of a brown pigment. *(C,G,K,O,S)* Helium ion micrograph of cross-sectioned scale. Each lamina is colorized, with approximate thickness indicated by an adjacent red bar (precise measurements were taken at sites chosen as in Methods). White scale bar is 500 nm and applies to all HIM images. *(D,H,L,P,T)* Reflection spectra for the adwing surface of disarticulated scales. Solid line is the mean spectrum, and blue envelope is one standard deviation; minimum N=6 spectra per graph.

**Figure 3:**
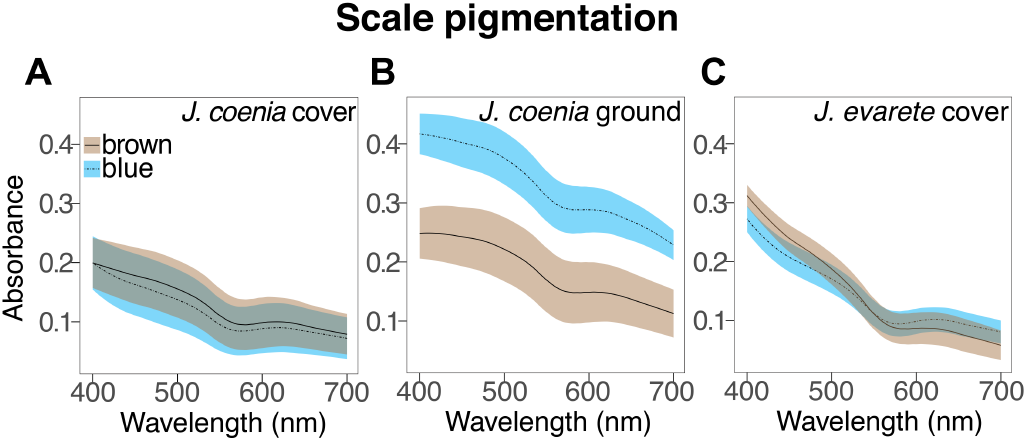
Absorbance spectra show the effect of artificial selection on scale pigmentation. *(A)* Absorbance measures in *J. coenia* wild-type (brown), and artificially selected (blue) cover scales show that both have comparable pigmentation (Mann-Whitney *U*, Table S1). *(B)* Absorbance of selected *J. coenia* ground scales is nearly doubled relative to brown wild-type scales. *(C)* Absorbance does not differ between blue and brown *J. evarete* cover scales, and is similar to pigmentation in *J. coenia* cover scales. Plots show mean spectra with envelope of one standard deviation, N=6 spectra per sample.

We next investigated whether ground scales also contributed to blueness after artificial selection. In artificially selected buckeyes, the ground scales generally had similar architecture to the cover scales, but with less uniform lamina color: ground scales exhibited a color gradient from the stalk outward (Fig. 4A-B’). Correspondingly, ground scales had a similar mean thickness but more variability than cover scales (190 ± 29 nm). Ground scales were much more heavily pigmented than cover scales (Fig. 3B, Fig. 4B”), such that the abwing surface was black (Fig. 4B). The extra pigmentation in ground scales enhances spectral purity by absorbing light transmitted through the cover scales, thus reducing backscatter and making the observed blue color more saturated (similar to [15]). We conclude that cover scale laminae are the major source of blueness in artificially selected buckeye butterflies, while melanic ground scales secondarily enhance spectral purity.

**Figure 4:**
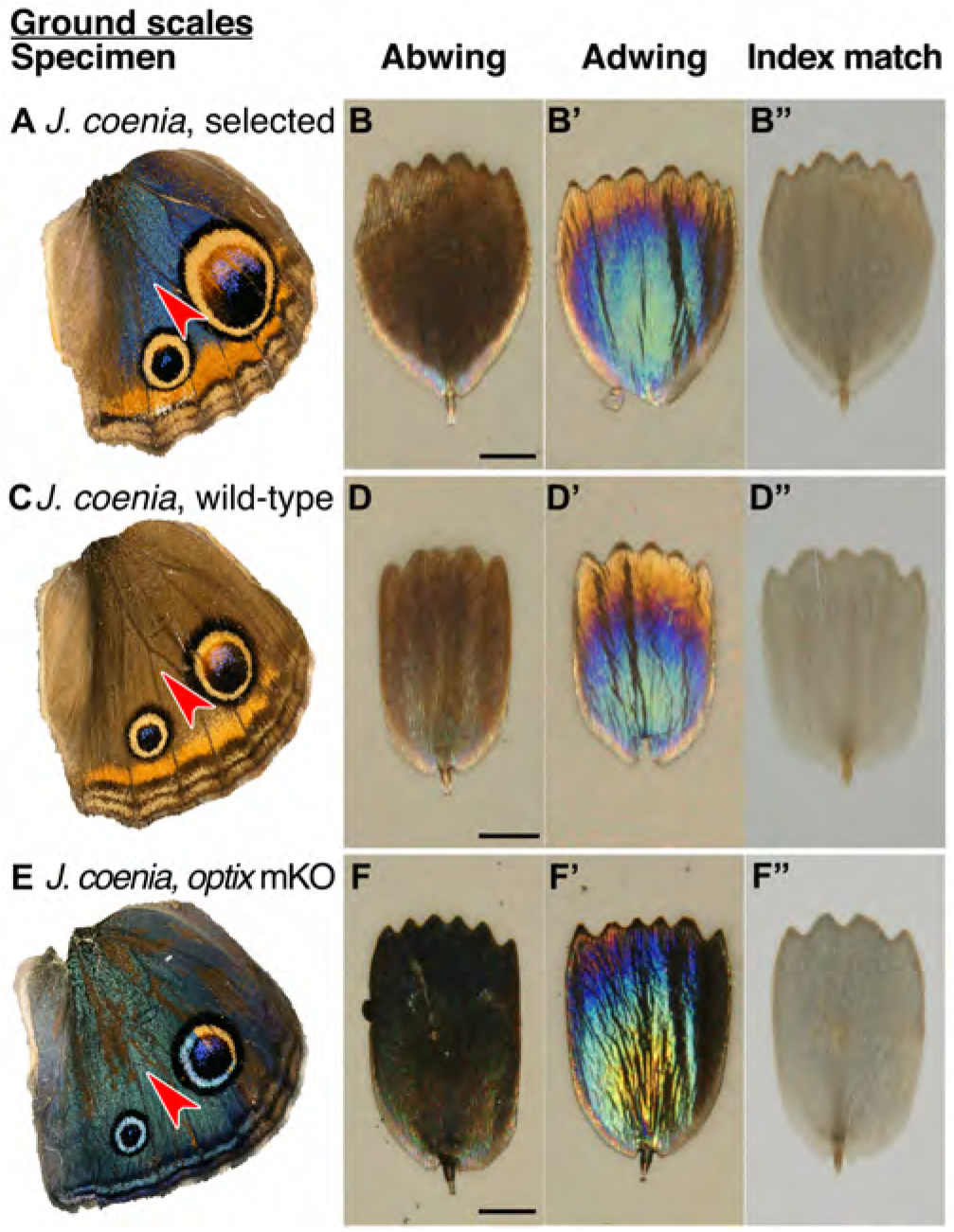
Structure and color of *J. coenia* ground scales. (*A,C,E*) Wings with red arrowhead indicating the region from which scales were sampled. (*B,D,F*) Scale in abwing orientation, i.e. ridges facing up. (*B’,D’,F’*) Scale in adwing orientation, i.e. lamina facing up. (*B”,D”,F”*) Scale immersed in fluid with a refractive index matched to chitin, thus eliminating reflection to show only pigmentary color. *(A-B”) J. coenia* artificially selected ground scales. *(C-D”) J. coenia* wild-type ground scales. *(E-F”) optix* mKO mutant ground scales. Scale bars are 25 µm.

For comparison, we tested the source of color in wild-type brown scales and found that they also had structural color (Fig. 2E-H). Brown cover scales had the same general architecture and no more brown pigment than did blue cover scales (Fig. 3A, Mann-Whitney *U*, Table S1). The salient difference was lamina thickness: brown scales were markedly thinner, measuring only 109 ± 12 nm (Analysis of Variance (ANOVA), p < 2×10^−16^, Fig. 1C, Fig. 2G). A 109 nm chitin thin film reflects a desaturated golden color due to reflectance of many long wavelengths. This golden structural color was confirmed by the adwing scale color, the color of the bare lamina in dissected scales, and the adwing reflectance spectra of brown scales (Fig. 2F’-F”, H). Therefore, though brown coloration is often attributed to pigmentation, wild-type brown cover scales also had a structural color, one simply tuned to enhance different wavelengths.

Artificial selection also altered the absorption and lamina thickness of the ground scales (Fig. 4A-D). The wild-type (brown) ground scales were thinner than the blue ground scales (151 ± 30 nm, ANOVA, p = 2×10^−6^, Fig. 1C). However, the mean difference was less extreme than in cover scales: blue cover scales were on average 78 nm thicker than wild-type, while blue ground scales were on average 39 nm thicker. Selected ground scales were markedly more absorbing than wild-type ground scales (Fig. 3B, Mann-Whitney *U*, Table S1), which is consistent with increased pigmentation that decreases backscatter in blue wing regions.

We conclude that the artificially selected buckeye butterflies rapidly evolved blue wing color via a 71% mean increase in lamina thickness in cover scales and a similar but less pronounced effect in ground scales. The effect was further amplified by increased pigmentation in ground scales, but without removing brown pigment from cover scales. Our results show that structural color can evolve quickly by modifying one dimension of an existing structure, and the process is facilitated by the initial presence of previously unrecognized structural color in wild-type brown *J. coenia*.

Since the artificially selected *J. coenia* wing pattern resembles natural iridescent variants in the sister species, *J. evarete* (Fig. 1F), we obtained hindwings of brown and blue *J. evarete* individuals from different geographic locations and tested whether blue cover scales in this species were also associated with increased lamina thickness (Fig. 2I-P). We found that the same mechanism explained color differences between geographic color variants: blue scales had 78% thicker scale laminae (blue 199 ± 14 nm; brown 112 ± 13 nm; ANOVA, p < 2×10^−16^, Fig. 1C) and no appreciable difference in pigmentation, compared to brown individuals (Fig. 3C, Mann-Whitney *U*, Table S1). Furthermore, in blue *J. evarete*, the ground scales were darkly pigmented. Thus, the artificially selected blue buckeyes faithfully recapitulate natural variation at the level of scale coloration between sister species.

### Color phenotypes in optix mutants include altered lamina thickness

Recently, Zhang *et al*. used CRISPR/Cas9 to generate mosaic knockout mutants of *optix* [16], a gene previously associated with pigment variation in butterfly wings [17]. Surprisingly, in addition to pigmentation phenotypes, *optix* mutants in *J. coenia* gained blue iridescence in wing scales. We tested phenotypically mutant blue scales from mosaic butterflies to determine what structural or pigmentary changes created the color change (Fig. 2Q-T). Where blue scales occured in the background region of the dorsal wing, blueness was due to similar factors as identified in artificially selected buckeyes. Lamina thickness of blue cover scales was substantially increased compared to wild-type brown scales (212 ±11 nm, ANOVA, p < 2×10^−16^, Fig. 1C). The concentration of brown pigment in the cover scales was significantly reduced relative to wild-type scales within the same mosaic wing (Fig 5A, Mann-Whitney *U*, Table S1) but comparable to selected animals (Fig. 3A, Table S1). Ground scales (Fig. 4E-F”) were likewise similar to those of selected blue animals, having thick and variable laminae (199 ±31 nm, ANOVA, p=5×10^−5^ versus wild-type. p=0.36 versus selected, Fig. 1C) and significantly increased pigmentation (Fig. 5B, Mann-Whitney *U*, Table S1). Overall, blue scale identity in *optix* mutants was caused by similar mechanisms as artificially selected blue.

**Figure 5:**
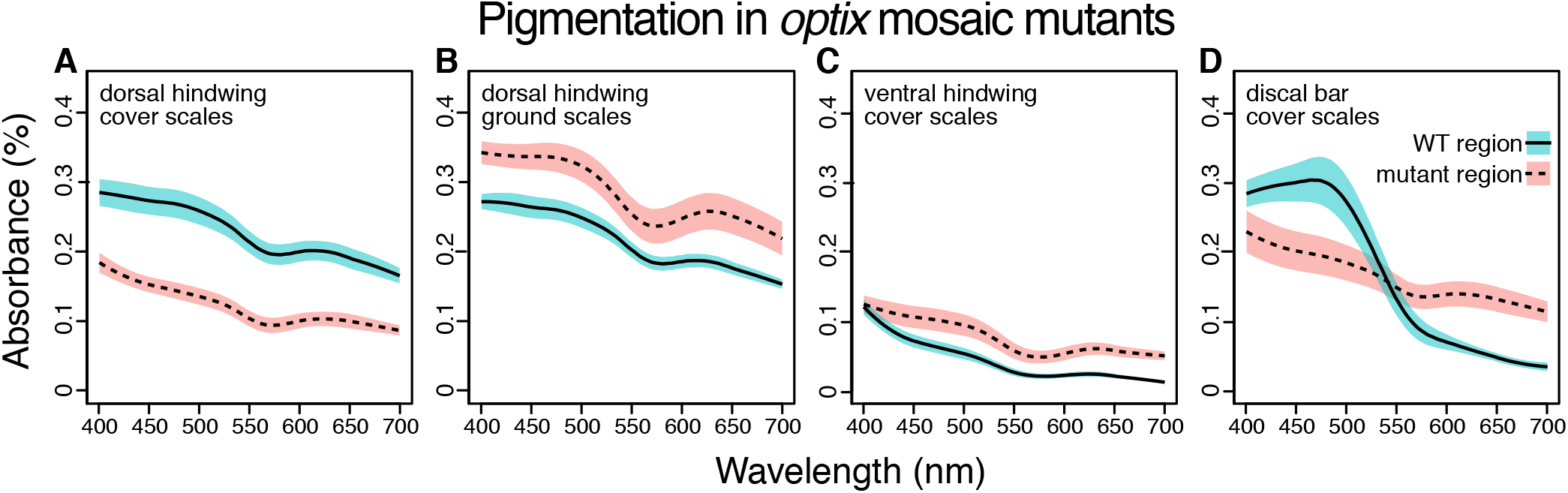
Absorbance spectra show the effect of *optix* knockout on scale pigmentation across wing pattern elements. All comparisons are between wild-type and mutant regions in the same mosaic wing. *(A) optix* mutation decreases absorption in cover scales from the main background region of the dorsal hindwing (Fig. 2Q). *(B)* Absorbance of ground scales from the dorsal hindwing (Fig. 4E) is increased in mutant scales. *(C)* Absorbance increases with *optix* mutation in ventral hindwing cover scales (Fig. 6A,C). *(D)* In the dorsal discal bars, (Fig. 6E,G) *optix* regulates a switch between orange and brown pigment. Plots show mean spectra with envelope of one standard deviation, N=6 spectra per sample. Differences for all four comparisons are statistically significant (Mann-Whitney *U*, Table S1).

*optix* mutant phenotypes also affected structural colors and pigments differently across wing pattern elements. As originally postulated [16], excess melanin was produced in some ventral wing regions (Fig. 6A-D, Fig. 5C). We also observed regions where both pigment and structure were dramatically changed. For example, discal bars on the dorsal forewing, which are normally orange, gained blue scales through both converting lamina structural color to blue and replacing orange with brown pigment (Fig. 6E-H, Fig. 5D). The kinds of pigmentation effects were diverse: *optix* mutation increased the quantity (Fig. 5B, C), decreased the quantity (Fig. 5A), or switched the identity (Fig. 5D) of the pigment in different scales (Mann-Whitney *U*, Table S1). Because the butterflies were mosaic mutants, some of this phenotypic variability could be due to genotypic differences between clones (i.e. mono-versus biallelic gene deletion, as well as the exact size of the deletion) [16]. However, much of the variation in outcome could also be observed within single clones that spanned multiple wing pattern elements (defined by the Nymphalid ground plan [18], Fig. S1), suggesting that the patterning roles of *optix* are quite context specific.

**Figure 6:**
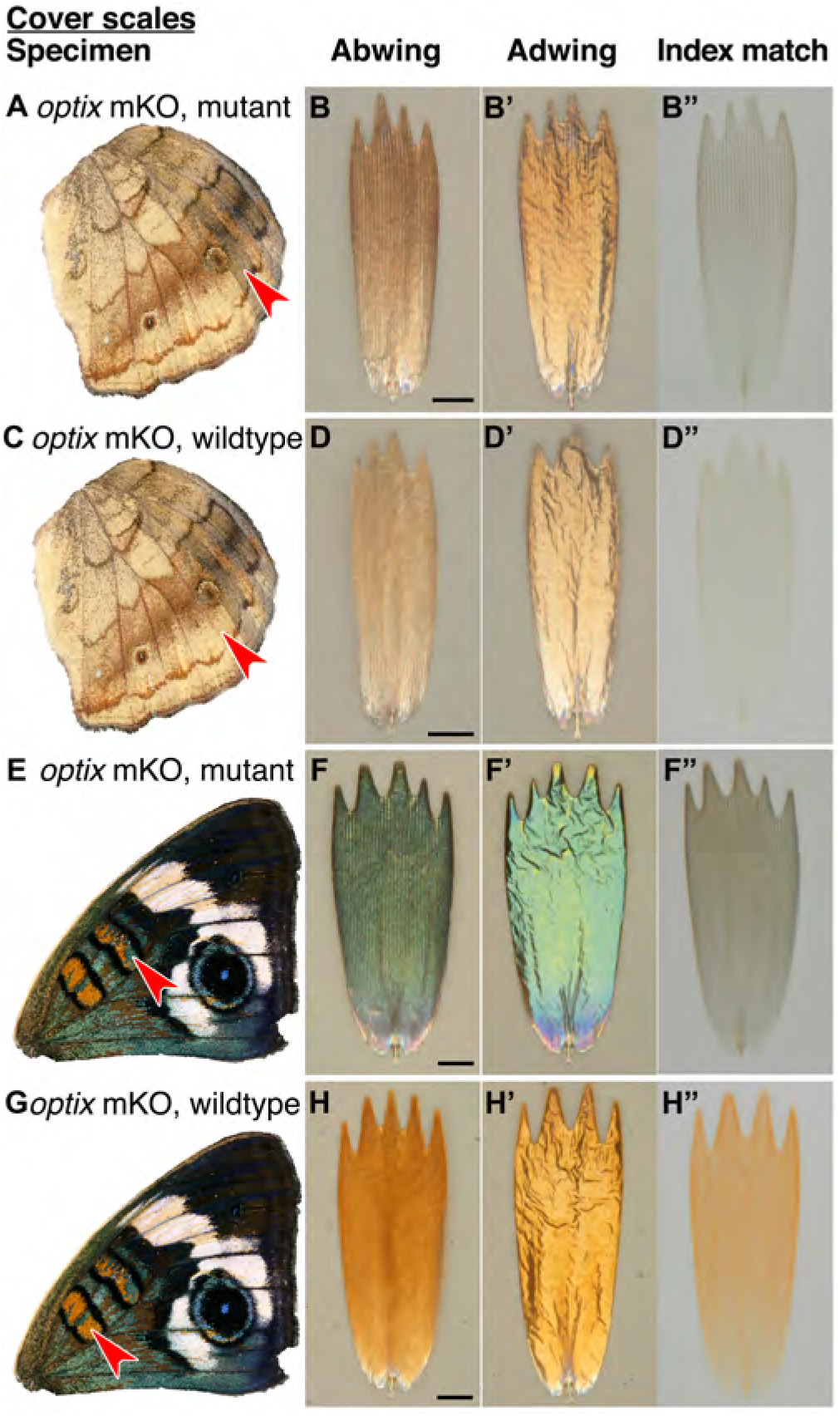
Effects of *optix* mutation on structure and color of *J. coenia* cover scales vary by wing region. (*A,C,E,G*) Wings with red arrowhead indicating the region from which scales were sampled. (*B,D,F,H*) Scale in abwing orientation. (*B’,D’,F’,H’*) Scale in adwing orientation. (*B”,D”,F”,H”*) Scale immersed in fluid with a refractive index matched to chitin to show only pigmentary color. *(A-B”)* Mutant cover scales from an *optix* mKO ventral hindwing have increased melanin. *(C-D”)* Wild-type cover scales from an *optix* mKO ventral hindwing. *(E-F”)* Mutant cover scales from an *opti*x mKO forewing discal bar have lost orange pigment, gained brown pigment, and increased lamina thickness, resulting in a shift to blue. *(G-H”)* Wild-type cover scales from the *optix* mKO forewing discal bar have both orange pigment and an orange lamina structural color. Scale bars are 25 µm.

In summary, *optix* knockout can have varied effects in a single scale by altering pigmentation, nanostructures, or both. These findings are consistent with *optix*’s described role as a developmental patterning gene that determines gross switches between discrete scale fates, and which, directly or indirectly, can regulate diverse downstream factors [19]. Since appropriate coloration critically depends on the proper combination of pigment and structural colors in both cover and ground scales (e.g. [20,21]), it is of particular interest that *optix* can regulate all of these components simultaneously. *optix* mosaic knockout mutants demonstrate that lamina thickness can be experimentally perturbed and highlight a multifunctional candidate genetic pathway for coordinated color evolution.

### Lamina thickness consistently predicts structural color wavelength

Relatives of *J. coenia* exhibit extensive color and pattern diversity, and blue structural colors in particular show patterns of variation that hint at ecological relevance (e.g. sexual dichromatism, seasonal polyphenism) (Fig. 7A). To assess the importance of lamina thickness variation in macroevolutionary color diversity, we sampled cover scales from nine species in the genus *Junonia* and a tenth species, *Precis octavia*, which belongs to the tribe Junoniini and exhibits seasonally polyphenic wing coloration. We prioritized large pattern elements that distinguish color forms within species. We compared scales using optical imaging, immersion index-matching, spectrophotometry, and Helium Ion Microscopy. All scales sampled had typical Nymphalid scale structure with a single plane of chitin forming the lower lamina.

**Figure 7:**
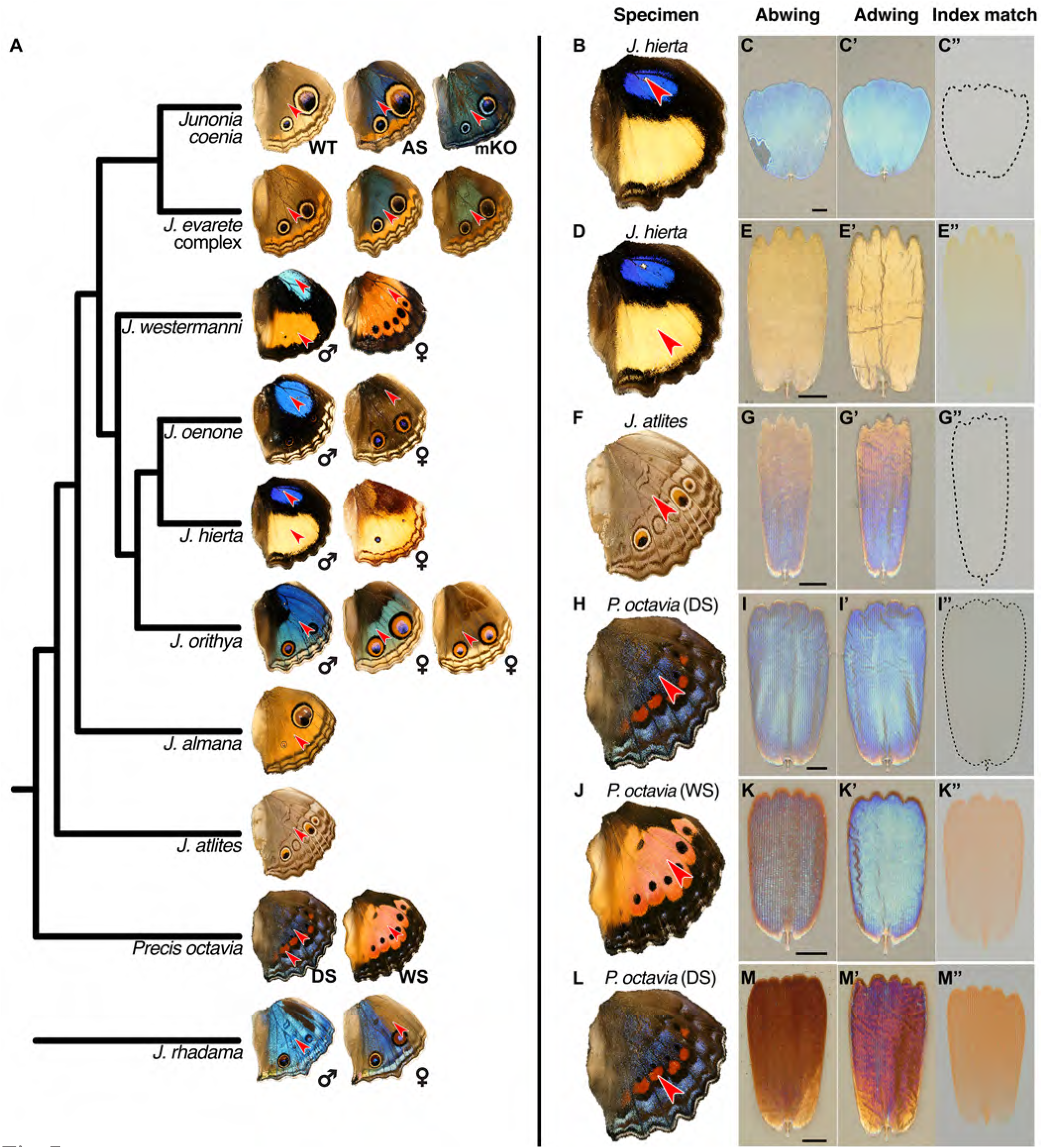
Lamina structural colors are an important component of overall wing color throughout *Junonia. (A)* Phylogeny of color variation in *Junonia* (based on [22]). Arrowheads indicate the color regions sampled for scale characterization. WT = wild-type. AS = artificial selection. mKO = *optix* mosaic knockout mutant. DS = winter/dry season form. WS = summer/wet season form. *J. evarete* variants are from different locations. Female *J. hierta* image is © Krushnamegh Kunte, NCBS. *(B,D,F,H,J,L)* Dorsal hindwing, arrow indicates the characterized scales’ location. *(C,E,G,I,K,M)* Abwing surface of cover scale. *(C’,E’,G’,I’,K’,M’)* Adwing surface of cover scale, showing lamina color. *(C”,E”,G”,I”,K”,M”)* Scale immersed in fluid with refractive index matched to chitin, thus showing only pigmentary color. *(B-C”) J. hierta* basal aura scales are unpigmented and appear blue due to lamina structural color. *(D-E”) J. hierta* has coordinated yellow pigment with a structurally yellow lamina. *(F-G”)* Neutral light grey of *J. atlites* is exclusively structural, due to additive color mixing of the multicolored lamina. *(H-I”)* Blue scales of dry season *P. octavia* are structurally colored since no pigment is present. *(J-K”)* Wet season *P. octavia* has discordant red pigment and blue lamina colors. The red pigment is localized in the ridges and cross-ribs on the abwing surface of the scale, while blue light from the lower lamina spills through the windows between them. *(L-M”)* The red band in dry season *P. octavia* is a more saturated red than in (J), due to the combination of both more red pigment and a structurally reddish lamina.

We tested whether the relationship between lamina thickness and color that we observed in experimental contexts applies more broadly. We sought to address two questions: First, does lamina thickness reliably predict lamina color, as measured from the adwing surface? While it is known that the thickness of a dielectric film controls the film’s reflectance, other variables such as refractive index, surface roughness, and pigmentation within the film also factor into reflectance, and these could plausibly vary among taxa. Second, how variable is lamina thickness? What range of thicknesses occur, and is there evidence for either quantized or continuous thickness variation? To address these questions, we measured reflectance spectra from the adwing surface of disarticulated cover scales from the 23 wing regions indicated in Fig. 7A. We then cross-sectioned scales, imaged with HIM, and measured thickness.

We found that lamina thickness varied continuously between 90-260 nm, indicating that all thicknesses over a more than 2.5-fold range are accessible (Fig. 8A). To better visualize the relationship between thickness and lamina color, we clustered similar samples into five color groups (Methods). Lamina colors in these groups could be described as gold, indigo, blue, and green, with a fifth variable group that included magenta, copper, and reddish colored scales (labeled as “red” in Fig. 8). Thickness differed significantly between all color group pairwise comparisons (Fig. 8A, ANOVA: p < 2×10^−16^, with *post hoc* Tukey’s Honestly Significant Difference test: p < 2×10^−6^ for all pairwise comparisons). The color groups were also associated with different reflectance profiles (Fig. 8B). In some cases, we obtained variable measures within individual specimens, which reflects biological color variation between adjacent scales, as well as varying color within individual scale laminae along their proximal-distal and lateral axes. A particularly striking example of the latter came from *J. atlites*. While the wing appeared light grey, at higher magnification individual scales could be seen to be multicolored (Fig. 7G’), and thickness measures from *J. atlites* overlapped the ranges of all color groups (Fig. 8A, see further analysis below).

**Figure 8:**
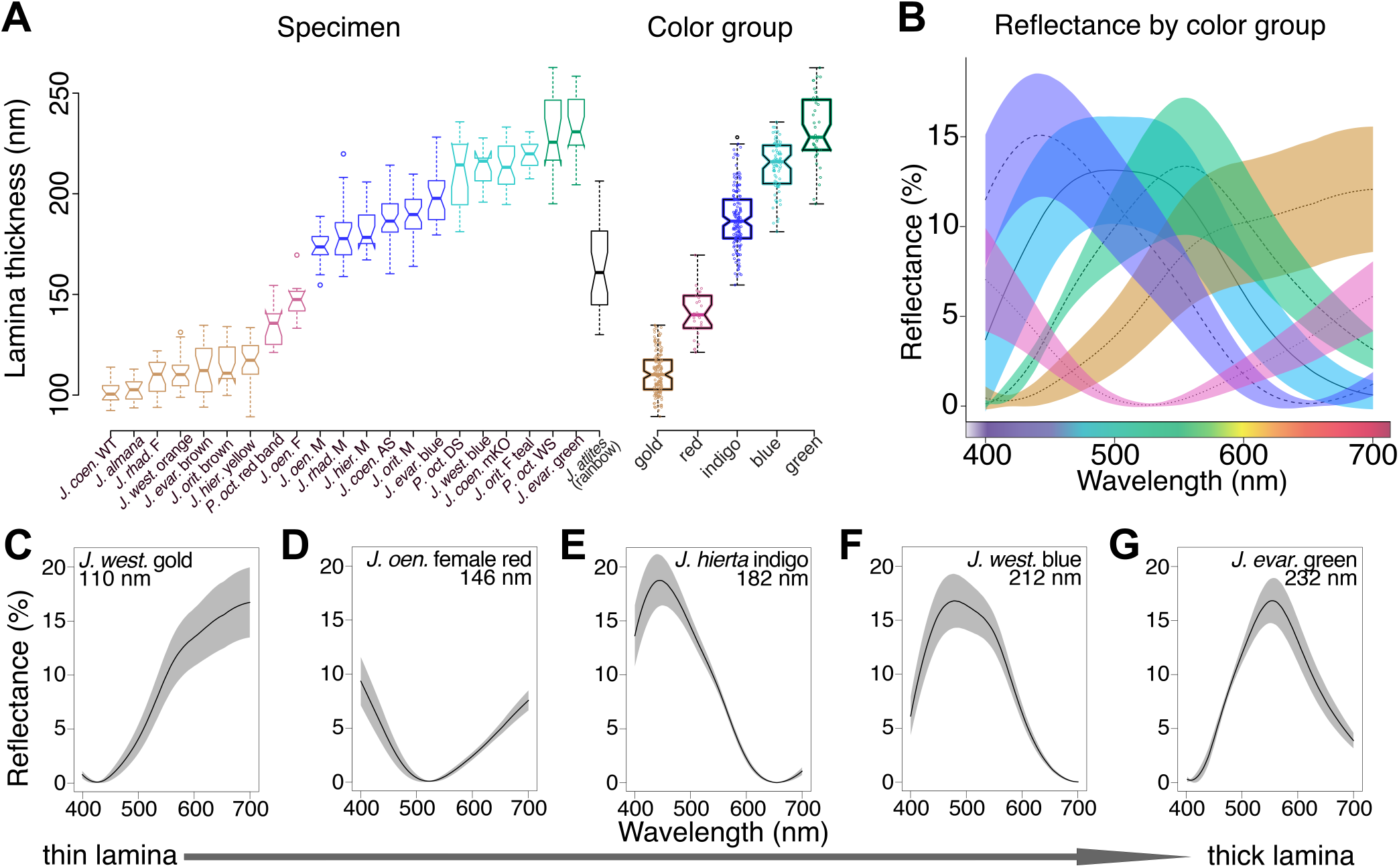
Lamina thickness predicts lamina color across the *Junonia* phylogeny. *(A)* Thickness measures for the regions indicated in Fig. 7A vary continuously over a 170 nm range (minimum N=3 scales and 12 measures per specimen). To visualize the relationship between thickness and color, we clustered similar specimens into five color groups described as gold, red, indigo, blue, and green. Thickness is significantly different between groups (ANOVA and Tukey’s HSD, p < 2×10^−6^). *J. atlites*, which has rainbow color gradients in each individual scale, has especially variable thickness, with measures overlapping the ranges of all color groups. Boxplots show median and inner quartiles, whiskers extend to 1.5 times the interquartile range, outliers are shown as points, and notches show 95% confidence interval of the median. *(B)* Color groups are associated with different reflectance profiles. Lines are mean spectra and envelopes show one standard deviation. N=6 spectra per specimen from panel A; clusters follow panel A. *(C-G)* Adwing reflectance spectra for representative individual specimens with increasing lamina thicknesses. The color sequence follows Newton’s series. Solid line is the mean spectrum and the envelope is one standard deviation; N=3 scales and 6 spectra per graph.

Lamina thickness had a consistent relationship with adwing scale reflectance for the taxa and color range we sampled. The order of color shift as lamina thickness increased followed Newton’s series, which is the characteristic color sequence for thin films [6,23]. This sequence can be understood in terms of an oscillating thin film reflectance function, which shifts toward longer wavelengths as film thickness increases (Fig. 8C-G). The thinnest films appeared gold due to reflectance of all the longer wavelengths (Fig. 8C). In mid-thickness laminae, a mix of two oscillations determined color: reflectance of the first oscillation was shifted toward far red wavelengths, while a second reflectance peak rose in the ultraviolet (Fig. 8D). Visible reflectance of thicker laminae was dominated by the peak of the second oscillation as it moved from indigo to green (Fig. 8E-G). That the trend between thickness and reflectance holds broadly suggests that color changes in *Junonia* butterfly scales have recurrently evolved via lamina thickness adjustments. Moreover, the consistency of the relationship between thickness and reflectance is useful. For example, structural variation could be rapidly surveyed by extracting fitted thickness estimates from reflectance measurements, a much less laborious process than sectioning for electron microscopy.

### Lamina structural color influences wing color throughout the genus Junonia

We next tested whether the extensive variation in lamina structural color among *Junonia* butterflies, explained by lamina thickness, also drives variation in overall wing color. An alternative hypothesis would be that composite wing color is usually dominated by pigmentation, particularly by pigments distributed on the outward-facing abwing surfaces of cover scales, above the lamina thin film. We measured pigmentation in cover scales from the same regions (Fig. 7A) to test the relative importance of pigments and lamina structural colors for wing color. (Structural colors and pigments are listed per each specimen in Table S2 and representative examples are shown in Fig. 7B-M”.)

Pigmentation was highly variable among *Junonia* species (Fig. 7B-M”, Fig. 9, Table S2). This included marked differences in pigmentation between regions of a single wing (e.g. yellow and blue regions in *J. hierta*, Fig. 7B-E”, 9A) and also variation between color forms and species throughout the genus (e.g. between sexes in *J. orithya*, Fig. 9C, and seasonal forms in *P. octavia* Fig. 7H-M”, 9B). Absorbance spectra varied in both shape and magnitude. Variation in magnitude, such as between the red band and the wet season morph of *P. octavia* (Fig. 9B), represents differences in pigment abundance. We also observed distinct absorbance spectral shapes, which can indicate the identity of the pigment (for example, contrast the spectral shape of the yellow pigment in *J. hierta*, Fig. 9A, versus red pigment in *P. octavia*, Fig. 9B, and brown pigment in *J. orithya*, Fig. 9C).

**Figure 9:**
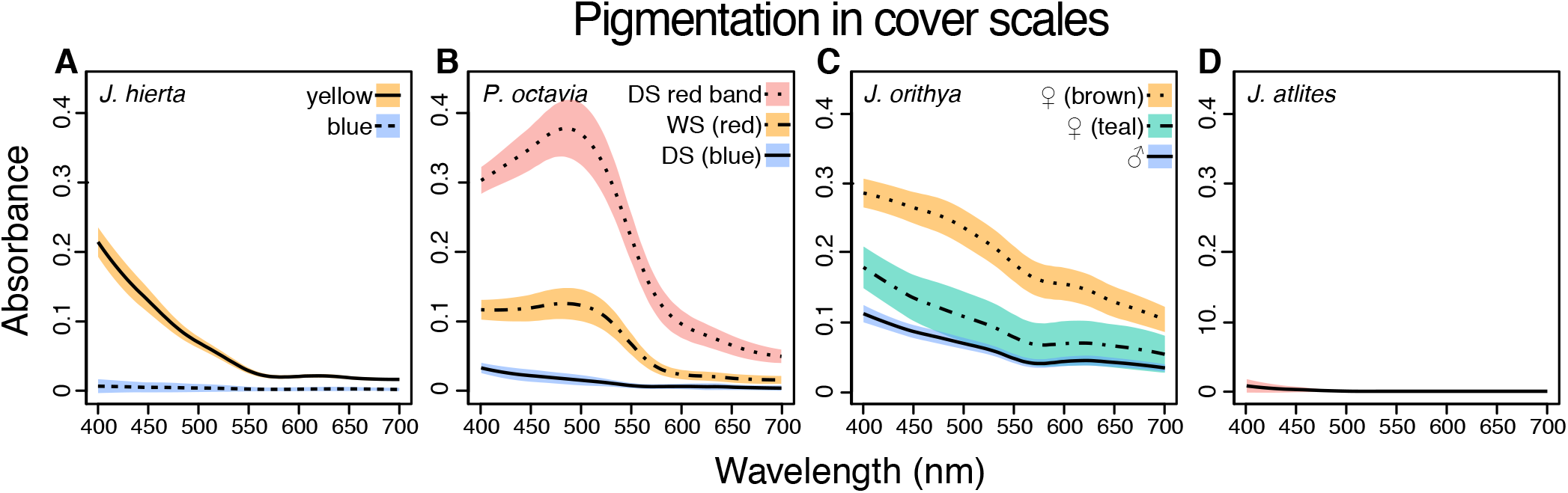
Absorbance spectra show variable pigment concentrations and identities among representative *Junonia* butterflies. Spectra were taken from cover scales from the regions shown in Fig. 7A. *(A) J. hierta* pigmentation varies by wing region (Fig. 7B-E”). *(B)* Extent of red pigmentation is the most important driver of color difference between seasonal morphs of *P. octavia* (Fig. 7H-M”). *(C)* Pigment absorbance differs by sex in *J. orithya*. *(D) J. atlites* scales lack pigmentation (Fig. 7F-G”). Plots show mean spectra with envelope of one standard deviation, minimum N=6 spectra per sample.

Notwithstanding the clear importance of pigmentation among *Junonia* butterflies, pigment variation was insufficient to explain the breadth of wing color diversity, and lamina structural colors made up the shortfall. The importance of lamina structural color was most obvious in scales that entirely lacked pigments. For example, the blue basal aura regions of male *J. westermanni*, *J. hierta*, and *J. oenone* wings had unpigmented cover scales with structurally blue laminae (Fig. 7B-C”, Fig. 9A). Most of the pigmentless scales we sampled were blue, with the notable exception of *J. atlites* scales (Fig. 7F-G”, Fig. 9D). These scales had rainbow gradient laminae, which presumably create the overall light grey by additive color mixing [24]. *J. atlites* demonstrates that lamina structural color can fundamentally drive wing color even in neutrally colored wing regions that are not obviously iridescent, and also that thickness can be patterned at fine spatial resolution within a single lamina.

In most wing regions, color was determined by the interaction of both lamina structure and pigments. For example, in the cover scales of *J. hierta* (Fig. 7D-E”, Fig. 9A), the yellow lamina structural color and yellow pigment were mutually reinforcing, with the lamina sensibly reflecting wavelengths that the pigment does not absorb. Other examples help delineate how much pigment is required to overpower the lamina color. In blue *J. evarete*, pigments in the cover scale ridges absorbed approximately 0.2 AU (Absorbance Units, i.e. 37% of light not transmitted, Fig. 3C) of the blue wavelengths that the lamina reflected most brightly (Fig. 2L). With this ratio, wing hue was still driven by the lamina structural color. The cover scales of *J. orithya* were similar (Fig. 9C), having a neutral dark pigment (i.e. a pigment that absorbs all visible wavelengths) in the scale ridges. Perhaps dark pigment in the ridges functions like a Venetian blind to limit iridescence, so that at high viewing angles, where iridescence would be most pronounced, light from the lamina is quenched.

Because of their range of pigment concentrations, *P. octavia* specimens were also useful to test the tradeoff between pigment abundance and lamina color influence. When viewed at high resolution, scales from the wet season morph of *P. octavia* contained red pigment in the ridges and ribs (max absorbance 0.12 ± 0.02, Fig. 7K,K’, Fig. 9B), while reflected light from the blue lamina spilled through the windows between ridges. Viewed macroscopically, this combination made a lightly saturated red. To display a richly saturated red, much more pigment was required, as seen in the red band of the dry season morph (max absorbance 0.38 ± 0.04, Fig. 9B, Fig. 7 L-M”). These reddest scales also had thinner, structurally magenta and copper colored laminae that may further reinforce redness (Fig. 8A, Fig. 7M’). The concentration of red pigment was the most important driver of the color difference between *P. octavia* seasonal morphs. The blue and red morphs had only a subtle difference in lamina thickness (Fig. 8A), and the laminae of both were blue (Fig. 7 I’, K’), but the blue morph lacked any red pigment (Fig. 9B).

Overall, *Junonia* wing color was determined by complex mix-and-matching of different lamina thicknesses and pigments. A thin film lower lamina was present in all scales, but its influence on wing color was adjusted by the amount and placement of pigment, especially in the upper surface of the scale. Pigments can mask lamina structural color at high enough density, depending on the placement and color of the pigment as well as the color of the lamina. In our tests, when pigmentation absorbed ≤ 0.2 AU of the relevant wavelengths, it did not cancel out lamina structural color.

### Comparison to thin film equation

We compared our empirical data to Fresnel’s classical thin film equations, which model the reflectance of an idealized dielectric thin film [7,25]. This model has previously been used to estimate the thickness of butterfly scale laminae based on their adwing reflectance spectra [8,21]. For each sample, we modeled the expected reflectance using our thickness measurements, and then compared to the measured reflectance spectra. We used 1.56 for the refractive index of chitin [26] and a maximal angle of illumination of 30° following [27] (because spectra were measured through an objective lens with a numerical aperture of 0.5). To account for measurement error, we modeled films over all thicknesses within one standard deviation of the measured mean per sample (red envelopes, Fig. S3 A). We also modeled films with Gaussian thickness distributions for each sample, following [15]. This model is analogous to a single uneven film with mean thickness and surface roughness defined by the measured thickness and sample standard deviation (solid red lines, Fig. S3 A).

We found that qualitatively the model describes the main behaviors of our data: reflectance oscillates with a given frequency and brightness, and the function shifts toward longer wavelengths as thickness increases. Quantitatively, mean maxima and minima in the reflectance function were offset laterally for every specimen, by about 40-80 nm, with the modeled curves blue-shifted relative to the observed. A similar blue shift has been reported in butterfly scale laminae before [9]. The comparison improves if we assume a higher refractive index or thickness. However, to align modeled and measured spectra would require either an impossibly high refractive index (around 1.75) or increased thickness outside the error range of our measures (20-25 nm thicker than mean measurements). Possibly the lateral offset is due to a combination of the former. Alternatively, these results could indicate that scales have additional properties not fully described by the model. There are a number of differences between the idealized film and real scales, including curvature of the film and possible birefringence of the ridges. The lamina itself may not necessarily have a uniform material composition or refractive index. For example, contrasting sublayers within the lamina (as in [28]) could create extra reflective interfaces. Thus, our data are compatible with the expected behaviors of thin films, but modeling the specific case of butterfly scale laminae with quantitative precision may require additional parameters or calibration to an empirical dataset.

## Discussion

This study leverages the simplest photonic nanostructures, thin films, to interrogate the evolution and genetic regulation of structural color in *Junonia* butterfly scales. While there is a large body of literature attributing optical properties to various biological nanostructures, such claims commonly rest on correlation between mathematical models and spectral measurements. Here, we use two different experimental manipulations of the structure (artificial selection on wing color and knockout of the *optix* gene) in addition to broad interspecies comparisons to establish that lower lamina thickness quantitatively controls structural color wavelength in *Junonia* butterfly scales. The relationship between lamina thickness and wavelength holds over a wide range of thicknesses (90-260 nm) that generate Newton’s color series for dielectric thin films. Moreover, lamina structural color is one important determinant of overall wing color, including in wing regions that also contain pigments. Lamina structural colors contribute to the color differences that distinguish sexes, species, seasonal variants, and selectively-bred lineages of *Junonia* butterflies, highlighting that quantitatively tuning lamina thickness is a vehicle for color evolution in both micro and macroevolutionary contexts.

Because the lower lamina is part of the typical architecture of butterfly scales, our findings have broad implications for future research on adult color in numerous butterfly taxa. Foundational literature drew a distinction between highly derived scales with vivid structural colors and “standard, undifferentiated scales,” which conform to the butterfly scale Bauplan, have a simple monolayer lower lamina, and “are not truly iridescent, i.e., they do not produce brilliant structural colors” [29]. However, within the past ten years, individual examples of thin film interference from the lower lamina have emerged in diverse Lepidoptera, including in simple scales [8,9,15,21,28,30,31]. These newer descriptions and our thorough examination of many scales indicate two points: first, although thin films are indeed less brilliant than some other classes of Lepidopteran photonic structures (thin films only reflect around 20% of incident light), they are a consequential source of structural color. Second, thin films occur in many butterfly and moth lineages and likely arose early in Lepidopteran evolution. The lower lamina has a thin film morphology in all scales that resemble the scale Bauplan, meaning that reflectance from the lamina is the shared condition except where it is masked by either heavy pigmentation or a derived structure with higher optical contrast. Because butterflies commonly produce multiple lamina colors across wing pattern elements and scale types, it is probable that the developmental genetic networks for quantitatively varying lamina thickness are deeply conserved as well. Hence, it will be useful to report which lamina colors are present, in addition to identifying pigments, when describing butterfly colors.

Physical constraints inherent to thin film colors may help explain the division of color space between pigments and photonic structures. It is not well understood why certain hues seem to be more often produced by pigments while others are more often produced by structural colors (e.g. the abundance of blue structural colors but lack of blue pigments in birds [32] and the rarity of one class of red structural color in birds and beetles [33]). In *Junonia*, we show that by tuning thickness, thin film laminae can produce nearly all the spectral colors (i.e. yellow, green, blue, indigo), and even light achromatic colors (e.g. light grey in *J. atlites*) via color mixing across a gradient. Yet thin films are fundamentally incapable of producing certain colors, notably dark brown, black, and pure red. The medium thickness films that most nearly approach red have inherently poor color properties due to the oscillating nature of the thin film reflectance function. Since the colors of mid-thickness films are a mix of two reflectance peaks (Fig. 8C), they are reddish but not pure or well-saturated, and are better described as copper, magenta, and purple. Further, mid-thickness films are not bright: they reflect less total visible light than other thicknesses we observed (compare Fig. 8D to 8C, E-G). By contrast, red, black, and brown are prevalent pigment colors in *Junonia*, making pigments and thin film structural colors complementary color palettes with little overlap. The optical limitations of thin films may have partially determined how pigment families and scale architecture evolved in early butterfly lineages, which in turn initialized whether pigments or structures provide the most accessible route to evolve specific hues during subsequent diversification.

Our findings uncover a link between artificially selectable responses in lamina thickness and natural butterfly color variation, and expand on a previous artificial selection study on butterfly wing color [9]. In both *J. coenia* and *B. anynana*, color shift was accomplished by modifying the dimension of an existing structure, the lower lamina, with pigmentation being less important. Since the selected taxa diverged 78 million years ago [34] this similarity may be informative about evolvability in nymphalid butterflies generally. However, artificial selection in *B. anynana* primarily increased thickness in the obscured layer of ground scales, which can only weakly influence color, whereas *Bicyclus* species with naturally evolved violet wing color have violet thin films in their cover scales. In our study, artificial selection continued longer (12 vs. 6 generations) and elicited a more extreme response (71% vs. 46% increase in lamina thickness). Moreover, in *J. coenia*, we show that lamina thickness increased in the cover scales and fully recapitulated the naturally evolved mechanism of structural color in the sister species *J. evarete*. The thickness increases caused a stark wing color change plainly visible by eye, with appropriate wing patterning that also resembled *J. evarete* (thickened blue scales filled the background dorsal wing, while eyespots, distal pattern elements, and the ventral wing were unaffected). Our results robustly connect a rapid microevolutionary process to macroevolutionary diversity.

By using butterflies with CRISPR/Cas9-generated knockout of the *optix* gene, we are able to provide insight into the genetic regulation of lamina thin films. It was previously known that the *optix* wing patterning gene can regulate a switch between wild-type brown and blue iridescent wing color in *J. coenia* [16], but the mechanistic basis for the color switch remained unknown. Specifically, it was unclear whether *optix* regulated scale structure itself, or whether *optix* deletion merely caused the loss of brown pigment, thus unveiling a pre-existing iridescent structure. Here, we show explicitly that in certain wing regions and scale types, *optix* deletion substantially increases lamina thickness. Our findings also amend the earlier conclusion that *optix* represses structural coloration in *J. coenia* [16]. Rather, by regulating lamina thickness, *optix* regulates the wavelength of a photonic structure that exists in both wild types and mutants. This distinction has implications for the likely identities and behavior of downstream genetic factors, as well as the developmental basis of mutant blue coloration. For example, rather than preventing a cascade of downstream genes from acting to erect a photonic structure *de novo*, *optix* may subtly regulate the expression of a gene or genes that directly regulate lamina thickness, such as chitin synthase. Additionally, we uncover disparate effects of *optix* deletion on pigmentation, including promoting, suppressing, and switching the identity of pigments in different scale types. In aggregate, these results show that *optix*’s functions in *J. coenia* are highly context specific, depending on both wing region and scale type (i.e. ground or cover scale). Moreover, because *optix* can regulate both pigmentary and structural color, the *optix* pathway is an especially interesting candidate for coordinated color evolution, and further work on the detailed regulation of *optix* and its downstream targets is called for.

In summary, thin film reflectors, a morphologically simple class of photonic structures, are experimentally manipulable and broadly employed in the lower lamina of *Junonia* butterfly wing scales. Lamina thickness explains variation in structural color wavelength, responds to selection on wing color, and is regulated by the *optix* wing patterning gene. Tuning lamina thickness facilitates both microevolutionary and macroevolutionary shifts in wing color patterning throughout the genus *Junonia*, making the buckeye butterflies a promising study system with which to decipher the genetic and developmental origins of structural color.

## Materials and Methods

### Butterfly specimens

Reared *J. coenia* were fed fresh *Plantago lanceolata* or artificial diet (Southland Products, Lake Village, AK) as larvae and kept at 27-30 °C on a 16/8 hour day/night cycle. Artificially selected blue *J. coenia* were purchased as larvae from Shady Oak Butterfly Farm in 2014 (Brooker, FL). Wild-type *J. coenia* were from an established laboratory colony, originally derived from females collected in Durham, North Carolina [35] (for the comparisons to both *optix* mutant and selected butterflies) or were collected in California (comparison to selected butterflies only). We acquired preserved specimens from various vendors and collaborators (Table S2). Species-level identification was generally unambiguous. However, relationships among Neotropical *Junonia* are not well-resolved and the limited molecular data available do not cleanly support current designations [36–38]. Two recognized species, *J. evarete* and *J. genoveva*, have large ranges with extensive overlap and many variable color forms, including both brown and blue. We therefore described three Neotropical specimens as belonging to the *J. evarete* species complex to avoid accidental misidentification. Available diagnostic details, including ventral antenna club color and full collection details, are in Table S2.

### Optical Imaging

Scales were laid on glass slides. Optical images of scales were taken with a Keyence VHX-5000 digital microscope (500-5000x lens). For refractive index matching, we used immersion oil (nD=1.56) from Cargille Laboratories (Cedar Grove, New Jersey), and imaged with transmitted light. Scales were dissected by hand using a capillary microinjection needle. Whole wings were also imaged on the Keyence VHX-5000, using the 20-200x lens.

### Microspectrophotometry

For reflectance spectra, individual scales were laid flat on a glass slide, with the adwing surface facing up. We collected spectra of the adwing surface with an Ocean Optics Flame-S-UV-Vis-Es spectrophotometer mounted on a Zeiss AxioPhot reflected light microscope with a 20x/0.5 objective and a halogen light source. Measurements were normalized to the reflectance of a diffuse white reference (BaSO_4_). Data were recorded with SpectraSuite 1.0 software with 3 scans to average and a boxcar width of 7 pixels. The software wizard determined optimal integration time from the reference sample; time was generally about .007 seconds. Spot size was roughly circular, 310 µm in diameter, and centered on the scale. We processed spectra in RStudio 1.0.153 with the package ‘pavo,’ version 0.5-4 [39]. We first smoothed the data using the *procspec* function with *fixneg* set to zero and *span* set to 0.3. We then normalized the data using the “minimum” option of the *procspec* function, which subtracts the minimum from each sample. Because we use a diffuse standard and scales are specular, raw spectra overestimate reflectance. We therefore followed [8] in dividing spectra by a correction factor. We used a smaller correction factor of only 2.5, because in our setup the scale does not fill the full field of view. Absorption spectra from scales submerged in index-matched oil were collected and processed similarly, but under transmitted light with an integration time of 0.01 seconds, and without the “minimum” option.

### Helium Ion Microscopy

Surface imaging by HIM provides increased depth of field and enhanced topographic contrast compared to Scanning Electron Microscopy for a range of biological and other materials [40], including butterfly wing scales [41]. Samples were prepared for HIM by laying the wing on a glass slide with the region of interest facing down, wetting with ethanol, and freezing with liquid nitrogen. We then promptly cross-sectioned the wing through the region of interest with a new razor blade. After the sample warmed and dried, we used a capillary microinjection needle to transfer individual cut scales onto carbon tape. Scales were placed overhanging the edge of a strip of carbon tape, with one end pressed into the tape. We optically imaged the tape strip as a color reference and then transferred the tape to the vertical edge of a 90° stepped pin stub (Ted Pella #16177). While non-conductive samples can be imaged by HIM using low energy electrons for charge neutralization, we found that the unsupported overhanging edges of our scales tended to bend due to local charging [42]. We thus sputter coated with 4.5 – 13 nm of Au-Pd using a Cressington 108auto or Pelco SC5. Images (secondary electron) of the sectioned scales were acquired with a Zeiss ORION NanoFab Helium Ion Microscope using a beam energy of 25 keV and beam current of 0.8 – 1.8 pA (10 µm aperture, spot size 4). We then used the line measurement tool in ImageJ software to measure lamina thickness from the micrographs. We corrected measurements for slight variations in working distance not accounted for by the software scale bar, using T_correct_ = (T_raw_)/9058 µm × *d* µm, where *d* is the measured working distance and 9058 µm is the reference working distance. Thickness of female *J. westermanni* scales was not measured because specimens were unavailable.

Even with vertical mounting, the sectioned surface of the scale was not always perfectly perpendicular to the direction of the imaging beam, largely due to the scales’ tendency to curve. Viewing angle is critical, since measurements taken from a projected image viewed under erroneous tilt could cause systematic underestimation of thickness. We therefore tilted the microscope stage until the scale lamina was perpendicular at the measurement site, as diagnosed by observing an inflection point in lamina curvature (i.e. a switch between the upper and lower surfaces being visible). Thickness was only measured at visible inflection points (Fig. S3 B-D). We performed a tilt calibration to test the precision of our inflection point criterion and determined that an inflection point was only visible if the sample was within 4-5° of perpendicular. Since erroneous tilt is limited to 5°, thickness underestimation is limited to 1 nm. Slight overestimations are likely, due to the sputter coating.

The sectioned scale shown in Fig. 1A was milled using the gallium ion beam of the Zeiss ORION NanoFab (beam energy 30 keV, beam current 300 pA).

### Analyses

Statistical analyses were conducted in R 3.2.2. For Fig. 8 A-B, specimens were grouped following the largest natural breaks in the data for two metrics, mean thickness and weighted average reflected wavelength, which were in good agreement.

### Modeling film thickness

We modeled the reflectance from chitin thin films as previously described [27], including integrating reflectance for values of θ from zero to the maximal angle of illumination (i.e. averaging reflectances to simulate the inverted cone of light collected by the objective lens used in microspectrophotometry, given its numerical aperture). Specifically, since our objective had NA=0.5, we calculated reflectance over values of θ from 0 to 30°, multiplied by 2πθ, and then averaged over the cumulative circular surface area. For the model with Gaussian thickness distributions, we followed [15] using n=400 observations from the simulated thickness distribution.

## Acknowledgements

We thank Linlin Zhang and Robert Reed for *optix* mutant wings; Karin van der Burg for *J. coenia* eggs; Masaki Iwata and Joji Otaki for *J. orithya* wings; and Krushnamegh Kunte for the image of female *J. hierta*. We thank Edith Smith for fantastic blue buckeyes, information about their origin, and the image in Fig. 1E. We are indebted to Ryan Null, Bodo Wilts, and Samuel Thayer for insightful discussions. Erika Anderson, Craig Miller, and Michael Nachman gave helpful feedback on the manuscript. Helium Ion Microscopy was performed at the Biomolecular Nanotechnology Center, a core facility of the California Institute for Quantitative Biosciences, University of California, Berkeley. Funding was provided by a National Science Foundation Doctoral Dissertation Improvement Grant DEB-1601815 (to R.C.T. and N.H.P.) and a National Science Foundation Graduate Research Fellowship DGE-1106400 (to R.C.T.).

