## Supplemental for "Structural color in *Junonia* butterflies evolves by tuning scale lamina thickness"

**This PDF file includes:**

Figs. S1 to S3
Table S1 to S2

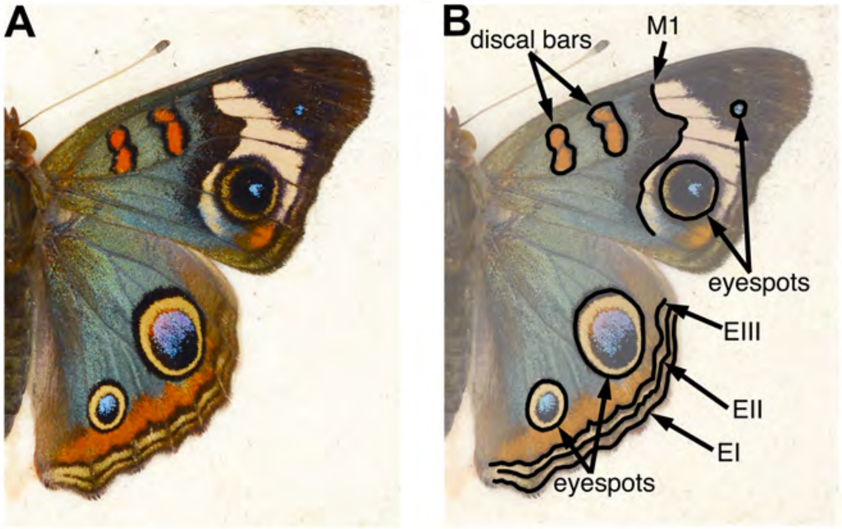

**Fig. S1.** *(A)* The artificially selected blue phenotype in *J. coenia* includes much of the dorsal wing surfaces. *(B)* Pattern elements of the nymphalid ground plan [18], which delineate the boundaries of blue regions, are labeled: discal bars, distal band of central symmetry system (M1), eyespots, parafocal element (EIII), and marginal and submarginal bands (EI, EII).

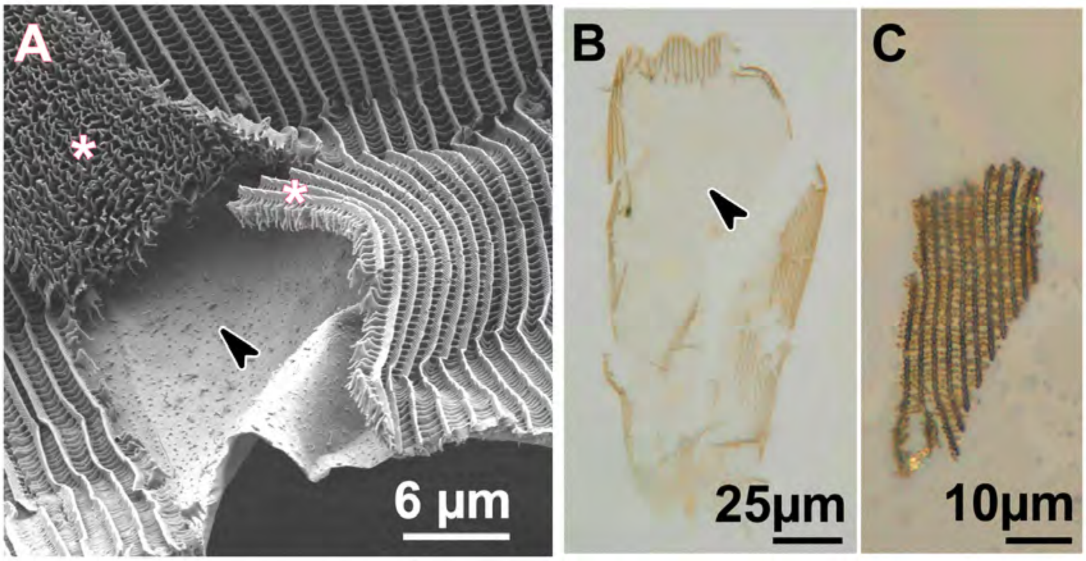

**Fig. S2.** Detailed characterization of dissected scales. *(A)* HIM image of a partially dissected scale. The arrow shows the exposed lamina and asterisks show detached swaths of all other scale components, i.e. ridges and cross-ribs. *(B)* Dissected cover scale from artificially selected *J. coenia* immersed in index-matched oil. Pigments are not primarily localized within the exposed lamina, shown by the arrowhead. *(C)* A swath of ridges and ribs removed intact from the scale in panel B. This piece of lamina-less scale is not blue.

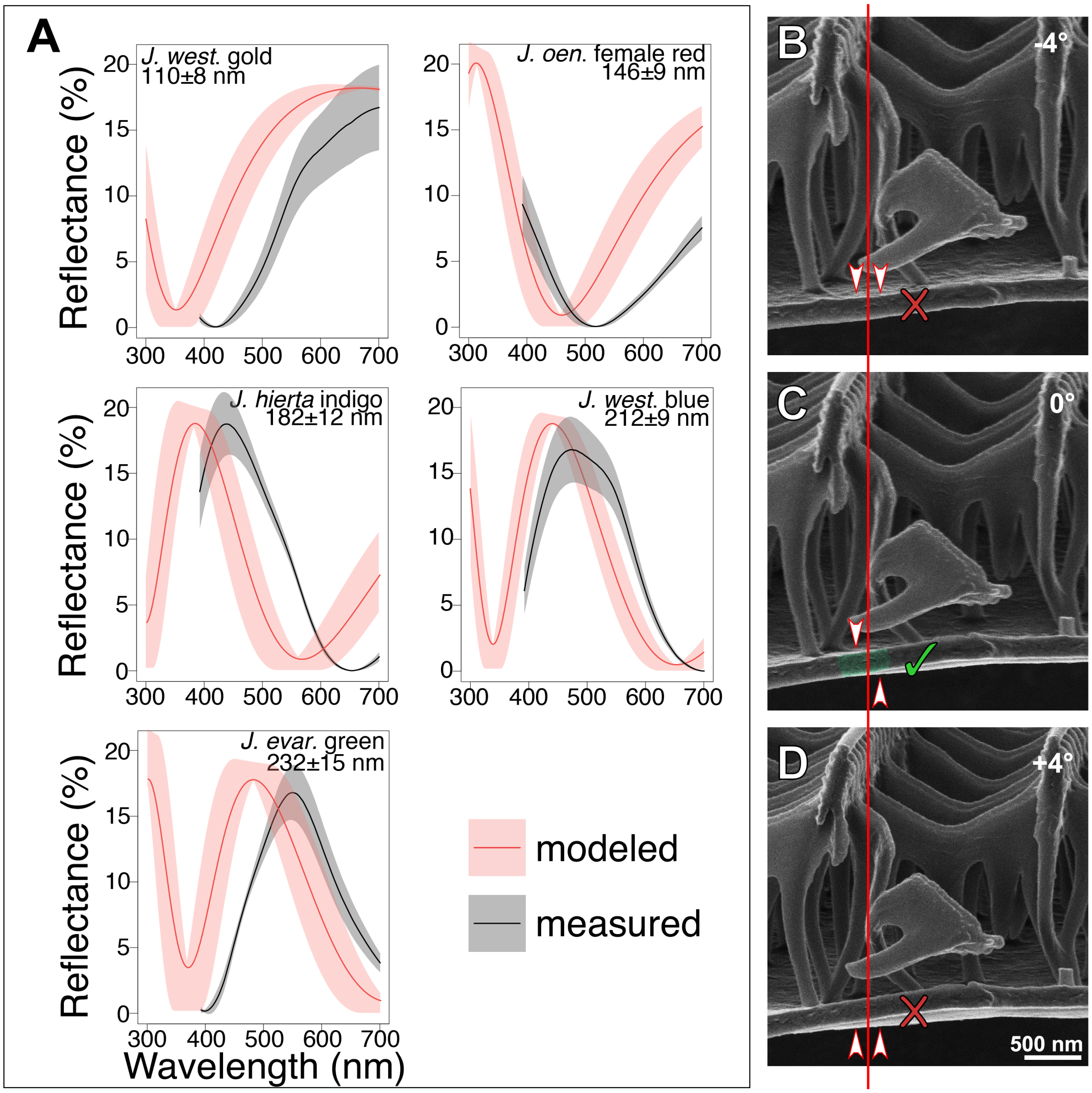

**Fig. S3.** *(A)* Modeled reflectance for representative specimens, using the classical Fresnel thin film equations, recovers the general shape of observed reflectance spectra. Black lines show the mean measured reflectance spectra; the grey envelope is one standard deviation (N=6 spectra per graph). In red, we show the modeled reflectance (solid red lines) and standard deviation (red envelopes), based on thickness measures from HIM images (See Methods). *(B-D)* Inflection point method for ensuring that lamina thicknesses are measured without sample tilt. At the position indicated by the vertical red line, an inflection point between the upper and lower surfaces of the lower lamina is visible only in (C). Arrowheads show whether the upper or lower surface of the lamina is visible in each panel, given the sample tilt and the lamina's curvature. If the lamina is tilted by 4-5° or more from the horizontal, no inflection point is observed at that position.

### *J. coenia:* wild-type and artificially selected

| nm | WT cover | AS cover | *U* | *p* |
|---|---|---|---|---|
| 400 | 0.199 | 0.200 | 35 | 0.9636 |
| 500 | 0.155 | 0.137 | 27 | 0.4371 |
| 600 | 0.098 | 0.088 | 27 | 0.4371 |
| 700 | 0.079 | 0.072 | 29 | 0.5532 |

| nm | WT ground | AS ground | *U* | *p* | |
|---|---|---|---|---|---|
| 400 | 0.248 | 0.417 | 72 | 0.0001 | ** |
| 500 | 0.221 | 0.376 | 72 | 0.0001 | ** |
| 600 | 0.149 | 0.288 | 72 | 0.0001 | ** |
| 700 | 0.113 | 0.228 | 72 | 0.0001 | ** |

### *J. evarete*

| nm | brown | blue | *U* | *p* | |
|---|---|---|---|---|---|
| 400 | 0.312 | 0.272 | 3 | 0.0082 | ** |
| 500 | 0.188 | 0.171 | 11 | 0.1807 | |
| 600 | 0.087 | 0.099 | 30 | 0.2343 | |
| 700 | 0.058 | 0.081 | 32 | 0.1375 | |

### *J. coenia: optix mutant vs artificially selected*

| nm | mut cover | AS cover | *U* | *p* |
|---|---|---|---|---|
| 400 | 0.184 | 0.200 | 23 | 0.4848 |
| 500 | 0.135 | 0.137 | 24 | 0.3939 |
| 600 | 0.100 | 0.088 | 20 | 0.8182 |
| 700 | 0.086 | 0.072 | 16 | 0.8182 |

| nm | mut ground | AS ground | *U* | *p* | |
|---|---|---|---|---|---|
| 400 | 0.342 | 0.417 | 36 | 0.0022 | ** |
| 500 | 0.323 | 0.376 | 32 | 0.0260 | * |
| 600 | 0.247 | 0.288 | 24 | 0.3939 | |
| 700 | 0.218 | 0.228 | 20 | 0.8182 | |

### *optix* mutants - regions of same mosaic wing

**dorsal hindwing cover scales**

| nm | WT | mut | *U* | *p* | |
|---|---|---|---|---|---|
| 400 | 0.285 | 0.184 | 36 | 0.0022 | ** |
| 500 | 0.259 | 0.135 | 36 | 0.0022 | ** |
| 600 | 0.200 | 0.100 | 36 | 0.0022 | ** |
| 700 | 0.165 | 0.086 | 36 | 0.0022 | ** |

**dorsal hindwing ground scales**

| nm | WT | mut | *U* | *p* | |
|---|---|---|---|---|---|
| 400 | 0.272 | 0.342 | 0 | 0.0022 | ** |
| 500 | 0.248 | 0.323 | 0 | 0.0022 | ** |
| 600 | 0.185 | 0.247 | 0 | 0.0022 | ** |
| 700 | 0.153 | 0.218 | 0 | 0.0022 | ** |

**ventral hindwing cover scales**

| nm | WT | mut | *U* | *p* | |
|---|---|---|---|---|---|
| 400 | 0.122 | 0.126 | 12 | 0.3939 | |
| 500 | 0.055 | 0.096 | 0 | 0.0022 | ** |
| 600 | 0.024 | 0.055 | 0 | 0.0022 | ** |
| 700 | 0.014 | 0.052 | 0 | 0.0022 | ** |

**dorsal forewing discal bar cover scales**

| nm | WT | mut | *U* | *p* | |
|---|---|---|---|---|---|
| 400 | 0.285 | 0.230 | 3 | 0.0152 | * |
| 500 | 0.272 | 0.185 | 0 | 0.0022 | ** |
| 600 | 0.072 | 0.140 | 36 | 0.0022 | ** |
| 700 | 0.036 | 0.115 | 36 | 0.0022 | ** |

**Table S1:** Comparison of absorbance values for individual scales immersed in refractive index matched oil. Mean absorbances at selected wavelengths are reported. Absorbances were compared using the Mann-Whitney *U* rank sum test. Minimum N=3 scales and 6 spectra per specimen. (* $p < 0.05$; ** $p < .01$) WT = wild-type. AS = artificial selection. mut = *optix* mosaic knockout mutant.

**Table S2. Butterfly specimens used in this study.**

| Genus | Species | Sex | Sample_origin | Other_diagnostics | SpecimenID_Fig5 | Color_cluster_Fig5 | Lamina color in cover scales | Pigment in cover scales |
|---|---|---|---|---|---|---|---|---|
| Junonia | almana | F | Mindoro; Philippines | | J. almana | gold | gold | orange |
| Junonia | atlites | M | Greater Sunda Islands; Sulawesi; Decmber 2006; Bioquip catalog #20B2293 | | J. atlites | NA (rainbow) | gradient of gold, copper, magenta, indigo, blue within each scale | generally none. the occasional scale has very slight brown |
| Junonia | coenia | M | Middle Island, Cool, California; May 2008; collector Chris Garcia | used as "wild-type" for comparisons to artificially-selected and in Fig.7,8 | J. coen. WT | gold | gold | dark brown, only in ridges + ribs |
| Junonia | coenia | M | reared; Rachel Thayer; UC Berkeley; ordered from Shady Oak butterfly farm (Brooker, FL, USA); October 2014; #00043 | | J. coen. AS | indigo | indigo | dark brown, only in ridges + ribs |
| Junonia | coenia | M | reared; Rachel Thayer; UC Berkeley; shipped as eggs from Karin van der Burg CornellU; July 2015; 126.41 | used as "wild-type" for comparisons to artificially-selected, and for thickness measures that were compared to optix mutants | J. coen. WT | gold | gold | dark brown, only in ridges + ribs |
| Junonia | coenia | M | reared; Linlin Zhang; CornellU; OptixKO-M30 | Used for Fig. 1,2,3,5 and discal bars in Fig. S2 | J. coen. mKO | blue | teal | dark brown, only in ridges + ribs |
| Junonia | coenia | M | reared; Linlin Zhang; CornellU; OptixKO-M61 | Used for ventral hindwing in Fig. S2 | NA | NA | very light white-gold to gold (wild-type regions). gold to copper (mutant regions) | dark brown, only in ridges + ribs. less in wild-type regions than mutant regions |
| Junonia | evarete complex | M | Coroico; Bolivia; Caranavi 11-12/2000; 700m; collector W. Zöller. | Labeled as Precis lavinia. Dark ventral antenna club. | J. evar. blue | indigo | indigo | dark brown, only in ridges + ribs |
| Junonia | evarete complex | M | Jamaica; Middlesex County; Manchester Parish; III-1980; Bioquip catalog #20B0320 | labeled as J. genoveva neildi. Light ventral antenna club. | J. evar. brown | gold | gold | dark brown, only in ridges + ribs |
| Junonia | evarete complex | M | unknown; collected prior to September 2014. | Light ventral antenna club | J. evar. green | green | green | dark brown, only in ridges + ribs |
| Junonia | hierta | M | Thailand (same specimen as yellow patch) | | J. hier. M | indigo | indigo | none |
| Junonia | hierta | M | Thailand (same specimen as blue patch) | | J. hier. yellow | gold | gold/yellow | yellow, more in distal part of scale |
| Precis | octavia | M | Tanzania; Tanga Region Amani Forest; October 2014 | dry season | P. oct. DS | indigo | variable between adjacent scales, ranges from indigo to teal | slight amount of brown, ridges+ribs only |
| Precis | octavia | M | Tanzania; Tanga Region Amani Forest; October 2014 | wet season | P. oct. WS | green | teal | red, only in ridges + ribs |
| Precis | octavia | M | reared ex ova; N. Bonkewitz | dry season | P. oct. red band | red | variable between scales and somewhat distal to proximal. Can be copper, magenta, or purple | lots of red |
| Junonia | oenone | F | Madagascar; Antananarivo province; Analamanga Region; V-2003; Bioquip catalog #20B0340 | labeled as P. oenone epiclelia | J. oen. F | red | variable; copper, magenta, or purple | lots of very dark brown/black |
| Junonia | oenone | M | Madagascar; Antananarivo province; Analamanga Region; V-2003; Bioquip catalog #20B0340 | labeled as P. oenone epiclelia | J. oen. M | indigo | indigo | none |
| Junonia | orithya | F | reared; Masaki Iwata; U of Ryukyus; No. 29; 08.12/1 ~ 12/9 | | J. orit. brown | gold | gold | dark brown, only in ridges + ribs |
| Junonia | orithya | F | Marinduque; Philippines | | J. orit. F teal | blue | teal | dark brown, only in ridges + ribs |
| Junonia | orithya | M | Malaysia | | J. orit. M | indigo | indigo | dark brown, only in ridges + ribs |
| Junonia | rhadama | F | Madagascar; February 2016 | | J. rhad. F | gold | gold | dark brown, only in ridges + ribs |
| Junonia | rhadama | M | Madagascar; January 2016 | | J. rhad. M | blue | blue | slight brown, only in ridges + ribs |
| Junonia | westermanni | M | Central Af Repub; Region de Bambiri; May 2016 | | J. west. blue | indigo | indigo | none |
| Junonia | westermanni | M | Central Republic Congo | | J. west. orange | gold | gold | orange |
| Junonia | westermanni | F | Central African Republic | | NA | NA | blue to indigo | red-orange |
